# Test-Retest Reliability of Dopaminergic fPET and fMRI Measures During Reward Processing

**DOI:** 10.1101/2025.11.03.686252

**Authors:** G Schlosser, PA Handschuh, M Murgaš, S Graf, C Milz, S Klug, P Falb, C Schmidt, B Eggerstorfer, E Briem, A Mayerweg, L Artmeier, GM Godbersen, L Nics, S Rasul, M Hacker, A Hahn, R Lanzenberger, MB Reed

## Abstract

Reward processing is essential to human brain function, with dopamine signalling in the nucleus accumbens (NAcc) as key element. The monetary incentive delay task is widely studied with functional magnetic resonance imaging (fMRI), measuring indirect hemodynamic changes. Functional positron emission tomography (fPET) with 6-[^18^F]FDOPA directly quantifies dopamine synthesis enabling dynamic assessment during task performance within a single scan. We investigated the reliability of 6-[^18^F]FDOPA fPET and blood oxygenation level dependent (BOLD) fMRI during a modified monetary incentive delay task in 25 healthy participants across two PET/MRI sessions. Intraclass correlation coefficients and coefficients of variance were computed for BOLD beta estimates and striatal dopamine synthesis at 30s and 2s resolutions. fPET showed fair to good reliability in the NAcc and putamen at rest, fair reliability during the win condition in the caudate and putamen, but poor reliability across the loss condition in all regions. Conversely, fMRI showed good reliability in the NAcc during feedback and in the caudate during feedback loss, but fair reliability elsewhere except for poor reliability in the caudate during cue loss. These findings indicate that both methods achieve comparable reliability but in different target areas, with the molecular specificity of fPET offering dynamic assessment of dopaminergic function.

**Clinicaltrials.gov Identifier:** NCT06675851

## Introduction

Dopamine plays a central role in modulating reward and loss processing (1). The dopaminergic neurons most relevant to these processes project from the ventral tegmental area (VTA) to the nucleus accumbens (NAcc) and ventral striatum (2,3). The NAcc is a abundantly innervated region important for the integration of cortical afferent information and goal-directed behaviour (4). Reward processing can be divided into an anticipation phase and an outcome phase, both of which engage dopaminergically innervated regions of the striatum, including the NAcc, caudate, and putamen (5). Disruptions in these processes are common in a variety of neuropsychiatric disorders including, major depressive disorders (MDD), addiction, anxiety and schizophrenia (6–8), where patients often show reduced motivation and anhedonia (9).

The monetary incentive delay (MID) task is a widely employed paradigm for probing the different phases of reward and loss processing in humans and has been combined extensively with blood oxygen level dependant (BOLD) functional magnetic resonance imaging (fMRI) (5). Originally based on non-human primate research showing reward anticipation-related activation in VTA dopaminergic neurons (1,10), the MID task has since been applied in more than 200 fMRI studies (5). BOLD fMRI has a number of advantages including a high spatiotemporal resolution, high sensitivity to fluctuations in blood oxygenation and the possibility to acquire high resolution anatomic scans in the same session for localisation and co-registration (11). However the BOLD signal is affected by a combination of multiple variables including blood oxygenation, blood flow and blood volume making it only and indirect measure of neuronal activation (12). Additionally the BOLD signal may be instable during longer task performance and can be influenced by physical effects like magnetic field inhomogeneities, low frequency drifts and heating (13,14).

Task-induced alterations during MID performance have recently been shown also for dopamine synthesis rates (15). Positron emission tomography (PET) with the radiolabelled dopamine precursor 6-[^18^F]FDOPA is a suitable method for mapping reward-related dopamine synthesis (16). However, conventional PET protocols typically require at least two measurements. One measurement is needed to capture resting state activity and another one captures task performance (17–19). The necessity to acquire data on different days introduces variability due to distinctions in daily performance and resting activity.

Furthermore, the radiation burden is increased by repeated measurements. To capture both resting-state and task-induced alterations in a single session functional PET (fPET) was developed to address these limitations (20,21). fPET allows to capture dynamics of radiotracer binding by controlling the delivery of the radiotracer to the blood via continuous infusion. Like fMRI, fPET uses repeated periods of task performance alternated with a control condition, thereby enabling the measurement of task-induced changes in dopamine synthesis across multiple conditions in a single scan session (15). To further optimize the fPET framework 6-[^18^F]FDOPA may be applied using a bolus plus constant infusion protocol, which allows for an increase in signal-to-noise ratio and provides a higher temporal resolution (22). As a further advancement, high-temporal resolution fPET with the glucose analogue [^18^F]fluorodesoxyglucose ([^18^F]FDG) and reconstruction to 3 s frames enabled a direct comparison between fPET and fMRI signals (23,24). Furthermore, [^18^F]FDG fPET showed high test-retest reliability during rest and moderate performance during the execution of a cognitive task, which was however still higher when compared to BOLD fMRI (25). For future scientific and clinical applications any imaging parameter must provide robust reliability to ensure detection of subtle task-dependent changes despite variance that may arise from repeated measurements. However, for 6-[^18^F]FDOPA dopamine synthesis fPET two crucial aspects for such use have not yet been determined: identification of task-induced changes at high temporal resolution of seconds and test-retest reliability. Therefore, we aim to assess the test-retest reliability of 2 s and 30 s frames of task induced dopamine synthesis measured with 6-[^18^F]FDOPA fPET and BOLD fMRI. We further aimed to assess the effects extended dynamic Non-local-Means (edNLM) filtering have on the test-retest reliability (26) of high-temporal fPET data compared to Gaussian 8 mm smoothing.

## Methods and Materials

### Subjects

Twenty-five (10 female, age 24.6 ± 6.1 years) healthy participants completed two PET/MRI measurements with the radiotracer 6-[^18^F]FDOPA. All subjects underwent a routine medical evaluation during a screening visit, which included electrocardiography, blood tests, neurological and physiological assessments, and a urine drug test. Female participants additionally took a pregnancy test at the screening visit and before each PET/MRI session. Psychiatric disorders were excluded using the Structured Clinical Interview for DSM-V, administered by an experienced interviewer. Exclusion criteria included a weight above 100 kg, current or past neurological, physiological, or psychiatric disorders, current breastfeeding or pregnancy, left-handedness, substance abuse, MRI contraindications, and participation in a study involving ionizing radiation exposure within the past 10 years. After detailed explanation of the study protocol, all subjects gave written informed consent. All subjects were insured and reimbursed for their participation. The study was approved by the Ethics Committee of the Medical University of Vienna (ethics number: 2321/2019) and all procedures were carried out in accordance with the Declaration of Helsinki. The study was registered in the European Clinical Trial Database (EudraCT 2019-004880-33).

### Monetary Incentive Delay Task

Participants performed a modified version of the monetary incentive delay (MID) task during simultaneous PET/MRI acquisition. They were instructed to maximize monetary gains and avoid losses by responding as quickly as possible to a target cue. Each trial began with the presentation of a cue indicating potential gain or loss amount. A response faster than the individualized reaction time (RT) resulted in the reward being obtained or the loss avoided; slower responses yielded no gain or incurred the loss. Reward/loss magnitudes were €0.5, €1, or €3, and participants started with an initial balance of €10. To enhance motivation, participants were informed that their remaining balance at the end of the session would be paid out if it was positive.

The task was adapted to accommodate the lower temporal resolution of fPET, which currently precludes conventional event-related analysis. Before scanning, participants completed a practice session to familiarize themselves with the paradigm and determine individual mean RT. During scanning, each participant completed four task blocks (297 s each), with each block containing 27 trials of fixed duration (11 s). Trials consisted of an anticipation phase (2–5 s, in 0.5 s increments), a reaction phase (limited to the individual RT; maximum 1 s), a feedback phase (2 s), and a baseline fixation cross of variable duration to maintain a total trial length of 11 s.

Blocks were assigned pseudorandomly as either gain or loss conditions. In gain blocks, the RT threshold was increased by 50 ms to enhance the probability of success; in loss blocks, it was decreased by 50 ms to increase the likelihood of failure. RTs were re-estimated at the start and midpoint of each block to adjust for fluctuations in participant attention. Between task blocks, participants were instructed to focus on a fixation cross and let their thoughts wander. Participants were informed regarding the RT threshold manipulation only after completion of both measurements. Full task design details are reported in Hahn et al. [17].

### PET/MRI data acquisition

Each subject underwent two 58-minute PET/MR scans using a hybrid PET/MR system (Siemens Biograph mMR, Erlangen, Germany at the Department of Radiology and Nuclear Medicine at the Medical University of Vienna). As dopamine synthesis rates depend on tyrosine plasma levels, participants fasted for at least four hours before each scan, avoiding sweetened beverages and caffeine. One hour before 6-[^18^F]FDOPA administration, participants received 150 mg Carbidopa and 400 mg Entacapone to inhibit peripheral metabolism of the radiotracer by dopa-decarboxylase and catechol-O-methyl transferase. Scans commenced simultaneously with the intravenous administration of 6-[^18^F]FDOPA. The radiotracer was administered using a bolus 816ml/h for 1 min and constant infusion 39 ml/h for 56 min, via a perfusion pump (Syramed µSP6000 with UniQUE MRI-shield, both Arcomed, Regensdorf, Switzerland).

The fPET data was acquired in list mode with the examination table moving in a stop-and-go pattern between brain and thorax field of view (FOV) as previously published by our group (22). This enabled non-invasive quantification of dopamine synthesis rates with a cardiac image-derived input function (IDIF). fPET acquisition started in the thorax FOV to obtain the IDIF from the left ventricle, ascending aorta, and descending aorta. After 6 minutes, the examination table was repositioned to capture the brain FOV, collecting baseline and MID task data (4⍰×⍰5 min) in a block design. Following each task block, the table moved back to the thorax to acquire additional IDIF data points (4⍰×⍰30s)(22). This alternating process was repeated several times to ensure the collection of consistent and accurate IDIF, baseline, and task data. fPET scans were obtained at a spatial resolution of (x, y, z) 2.09⍰×⍰2.09⍰×⍰2.03 mm and a matrix size of 344⍰×⍰344⍰×⍰127 voxels.

Structural MRI of the brain was acquired before fPET using a T1-weighted MPRAGE sequence (TE/TR⍰=⍰4.21/2200 ms, TI⍰=⍰900 ms, flip angle⍰=⍰9°, matrix size⍰=⍰240⍰×⍰256, 160 slices, voxel size⍰=⍰1 mm isotropic, TA⍰=⍰7:41 min) for spatial normalization.

The thorax was imaged using a T1-weighted STARVIBE sequence (TE/TR⍰=⍰1.44/3050 ms, flip angle⍰=⍰5°, matrix size⍰=⍰320⍰×⍰320, 208 slices, voxel size⍰=⍰1.19⍰×⍰1.19⍰×⍰1.2 mm, TA⍰=⍰5:33 min) for IDIF extraction.

MID task functional data were acquired using an echo planar imaging (EPI) sequence (TE/TR⍰=⍰30/2000 ms, flip angle⍰=⍰90°, matrix size⍰=⍰80⍰×⍰80, 34 slices, voxel size⍰=⍰2.5⍰×⍰2.5⍰×⍰2.5 mm with a 0.825 mm gap).

### Image derived input function (IDIF)

The MRI T1 STARVIBE, MRAC and mean PET images were used as templates to manually place three volumes of interest (VOIs) with fixed sizes on the left ventricle, the ascending aorta and descending thoracic aorta. Activity in these VOIs was sampled during our measurement protocol at 5, 18.5, 30, 41.5 and 55.5 min after start of tracer application. Afterwards, mean activity was extracted from each VOI across time points to derive IDIFs. Composite IDIFs were obtained by using the initial peak activity in the left ventricle VOI, as shown to be robust in prior studies (22), and by linear fitting the time activity curve in each VOI using 15 time points per subject to reduce motion sensitivity. Accuracy of the fit was further improved by incorporating venous blood samples collected at 7, 20.5, 32, 43.5 and 51.5 minutes, which were temporally aligned with PET frame acquisition via linear interpolation. The resulting IDIFs were then multiplied with the plasma-to-whole-blood ratio. The latter was fitted as a linear function. For more information on IDIF acquisition, see Reed et. al. (22).

### fPET data preprocessing

fPET list mode data was reconstructed with a Poisson ordered subset expectation maximization algorithm for each block (3 iterations, 21 subsets). Concatenation and decay correction of thorax and brain frames was performed separately at the start of the measurement. To adjust for the initial tracer kinetics the first thorax block was binned to 20×5s, 8×10s and 5×30s frames. The later thorax acquisitions were binned to 30s frames. The brain acquisition frames were binned into 30 s or 2 s frames, respectively. Attenuation and scatter correction for the brain scans was performed based on the structural T1 scans in a pseudo-CT approach (27). The thorax blocks on the other hand were corrected utilizing a DIXON MRAC with CAIPIRINHA sampling pattern (28).

fPET data was pre-processed as published previously by our team using SPM12 (Welcome Trust Centre for Neuroimaging) (29). Brain blocks were corrected for head movement (quality = best, registered to mean). After coregistration to the structural T1 scan, the images were normalized to MNI space. Furthermore, both 30s and 2s frames were smoothed with an edNLM Filter (26) with a kernel size of 3^3^ x 5 frames or with a conventional 8mm Gaussian kernel. A mask was applied to only include grey matter voxels.

To distinguish baseline metabolism and task dependant effects the fPET toolbox was utilized (30). In short, a general linear model was used, including regressors for win- and loss-blocks. For further optimization, the principal components of the six motion parameters which explained the majority of variance, were added as motion regressors. The baseline metabolism was defined as the average activity across grey matter voxels, that were not active during fMRI task performance (contrast success > failure, p < 0.001 uncorrected) and had not been identified previously as active voxels in a MID-task meta-analysis (31). The single frame before and after table movement was deweighted to 0.5 to reduce potential movement artifacts and the infusion start was anchored within the GLM by introducing a value of 0 kBq/cm^3^ at time 0. The net influx constant (Ki) was estimated using the Gjedde-Patlak plot with the slope fitted from t*=30 min after infusion start.

### BOLD signal changes

fMRI data was pre-processed as previously published by our team in SPM12 (32). We carried out slice timing correction to the middle slice and realignment to the mean image (quality = 1). Subsequently, BOLD data was normalized to MNI space and smoothed using an 8mm Gaussian kernel. First level analysis in the general linear model was conducted in an event-related design with one regressor for each event (cue win, cue loss, feedback win and feedback loss) and further regressors for head motion, white matter and cerebrospinal fluid. Afterwards four contrasts of interest were calculated from the GLM’s beta values: cue win vs rest, cue loss vs rest, feedback win vs rest, and feedback loss vs rest.

### Region of interest definition

Regions of interest (ROIs) were defined based on our previous 6-[^18^F]FDOPA fPET studies and a large scale meta-analysis of fMRI studies utilising the MID task (5,15). Regions of interest chosen for this analysis were the caudate, putamen and NAcc, extracted from the Harvard-Oxford atlas (33), see figure 2. f

**Figure 1:**
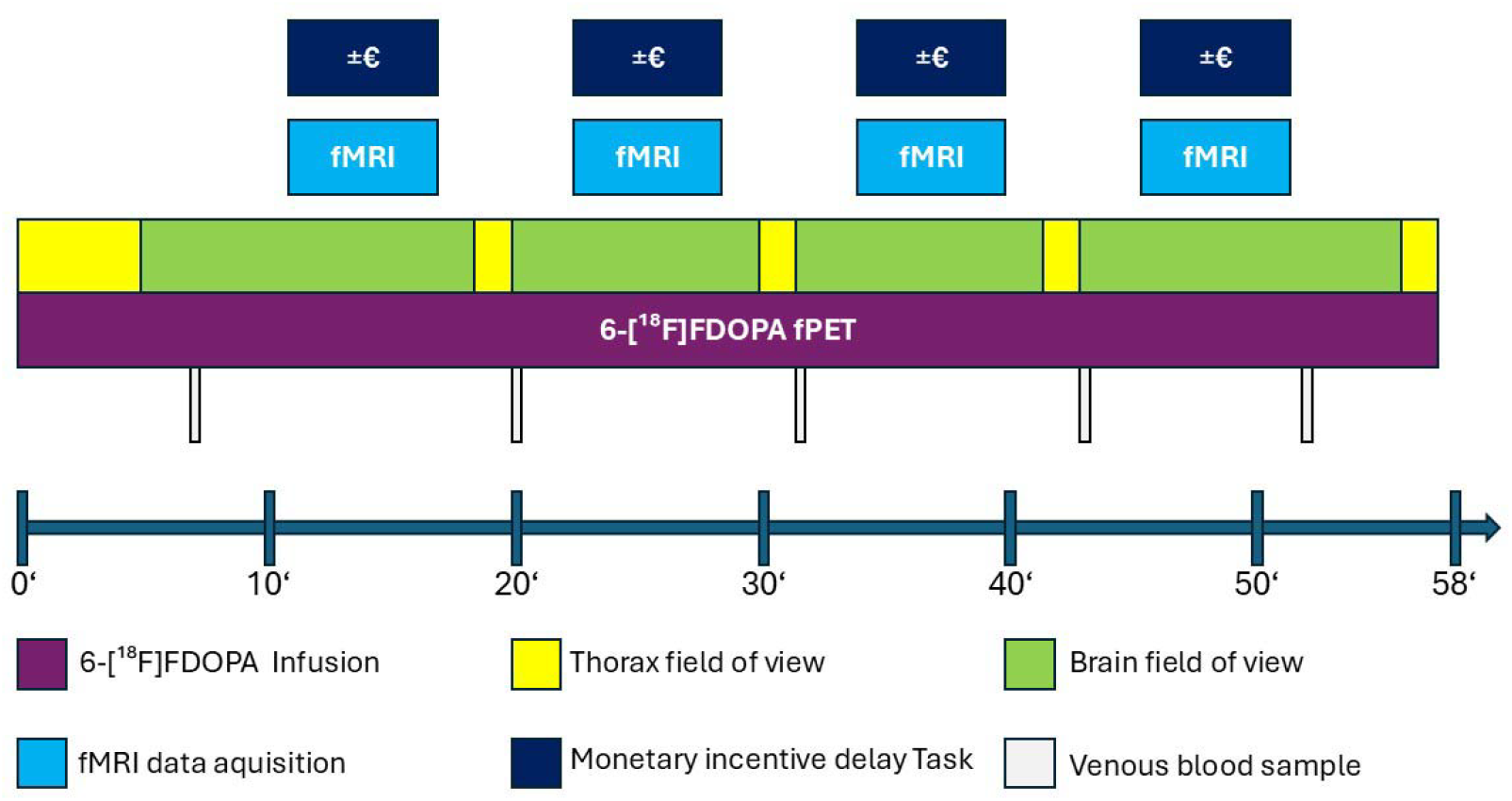
Graphical overview of the fPET/fMRI measurements. The fPET data acquisition always commenced with the PET field of view (FOV) positioned over the thorax (yellow box) for 6 minutes to capture the initial tracer peak. Afterwards, the table position was then shifted to the brain (green box), where fPET acquisition was initiated while participants fixated on a cross and engaged in unconstrained thought. At minute 11 after infusion start, the first MID task block began concurrently with the onset of the fMRI sequence. Manual venous blood samples were obtained at minutes 7, 20.5, 32, 43.5 and 51.5. Upon completion of the brain acquisition, the bed automatically returned to the thorax to collect additional data points for the image-derived input function (IDIF) using a stop-and-go motion protocol. This cycle was repeated multiple times to ensure reliable estimation of both IDIF and fPET task-related measures.

**Figure 2:**
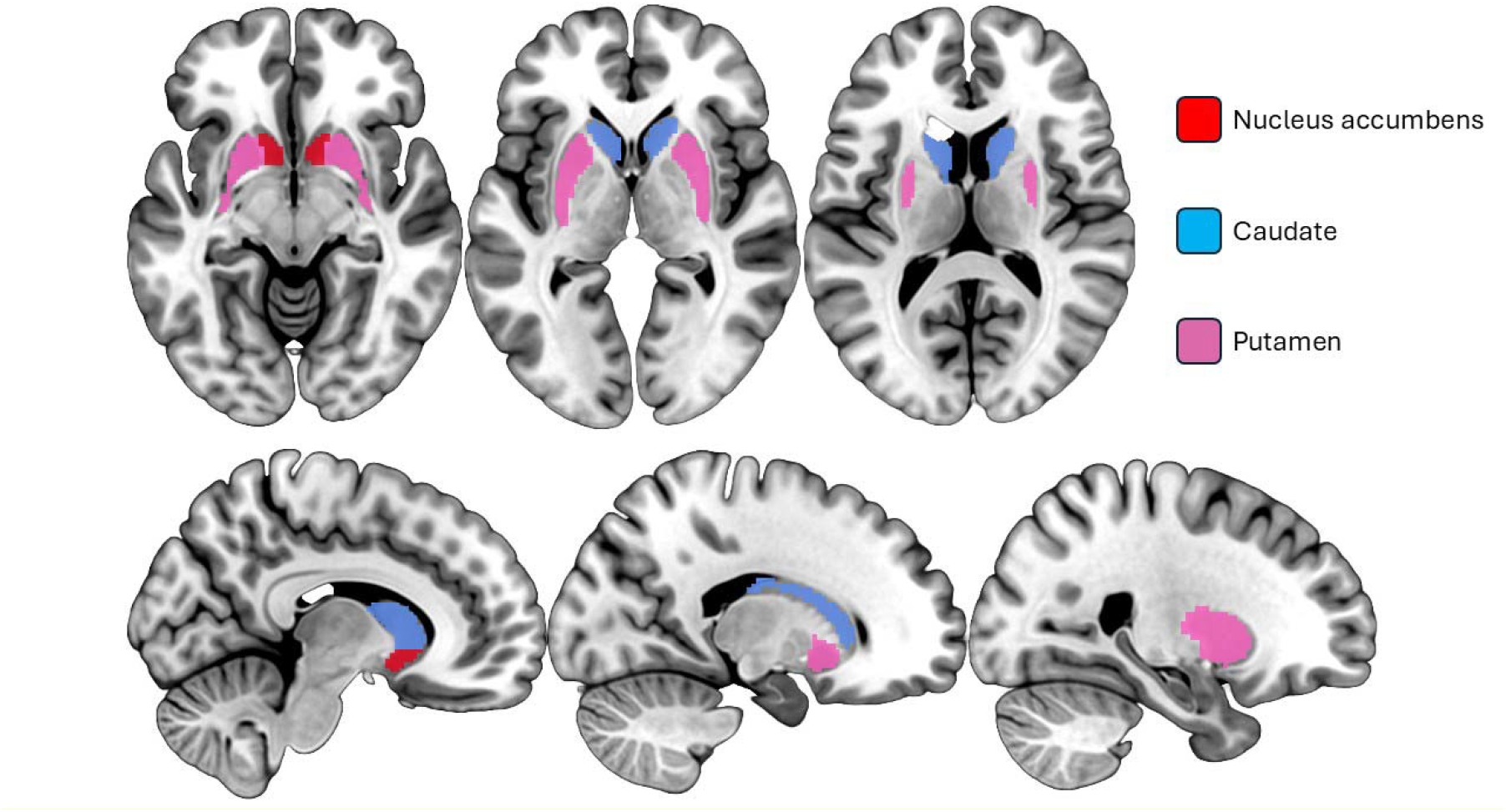
Regions of Interest including the nucleus accumbens (red), putamen (pink) and caudate (blue) in coronal and sagittal view, extracted from the Harvard-Oxford atlas (33).

### Statistical analysis

For 6-[^18^F]FDOPA fPET, the dopamine synthesis rate was estimated by calculating the net influx constant (Ki) to allow comparison with previously published studies. Similarly, fMRI beta values of the four different contrasts were computed for the same purpose. To quantitatively assess the consistency between the two PET/MRI measurements, the intraclass correlation coefficient (ICC, Equation 1) and the coefficient of variation (CV) was calculated for each modality, region of interest (ROI), and condition.

## Results

### fPET dopamine synthesis

For 30 s fPET data after edNLM filtering (see Tables 1 & 2), fair reliability was observed at rest in the NAcc (ICC = 0.58; CV = 3.13) and the putamen (ICC = 0.47; CV 3.86). Furthermore, fair reliability for the 30 s fPET data was observed during the win condition in the caudate (ICC = 0.54; CV = 23.97) and putamen (ICC = 0.57; CV = 19.24). In contrast, poor reliability was found across all regions during the loss condition (ICC = 0.30-0.37; CV = 29.83-37.53), in the NAcc during the win condition (ICC = 0.32; CV = 34.01), and in the caudate at rest (ICC = 0.39; CV = 7.20).

**Table 1:** Intraclass correlation coefficient (ICC) Values for fMRI beta values and fPET data for each region of interest and condition.

| ICC |  | Caudate | Putamen | N. accumbens |
| --- | --- | --- | --- | --- |
| fMRI | Cue win | 0.41 | 0.45 | 0.46 |
|  | Feedback win | 0.56 | 0.44 | 0.65 |
|  | Cue loss | 0.26 | 0.40 | 0.46 |
|  | Feedback loss | 0.61 | 0.44 | 0.67 |
| 30s edNLM fPET | Rest | 0.39 | 0.47 | 0.58 |
|  | Win | 0.54 | 0.57 | 0.32 |
|  | Loss | 0.37 | 0.30 | 0.31 |
| 2s edNLM fPET | Rest | 0.39 | 0.54 | 0.66 |
|  | Win | 0.57 | 0.46 | 0.29 |
|  | Loss | 0.33 | 0.26 | 0.11 |
| 2 s 8 mm Gaussian fPET | Rest | 0.38 | 0.51 | 0.57 |
|  | Win | 0.54 | 0.47 | 0.27 |
|  | Loss | 0.34 | 0.26 | 0.08 |

**Table 2:** Within-subject coefficient of variation (CV) Values for fMRI and fPET data for each region of interest and condition.

| CV |  | Caudate | Putamen | N. accumbens |
| --- | --- | --- | --- | --- |
| fMRI | Cue win | 76.70 | 47.26 | 75.76 |
|  | Feedback win | -143.33 | 379.33 | 88.17 |
|  | Cue loss | 77.22 | 46.48 | 82.74 |
|  | Feedback loss | 134.15 | 668.63 | 244.48 |
| 30s edNLM fPET | Rest | 7.20 | 3.86 | 3.13 |
|  | Win | 23.97 | 19.24 | 34.01 |
|  | Loss | 29.83 | 32.38 | 37.53 |
| **2s edNLM fPET** | **Rest** | 5.75 | 3.79 | 3.30 |
|  | **Win** | 23.04 | 23.75 | 35.57 |
|  | **Loss** | 28.88 | 33.17 | 40.55 |
| **2 s 8 mm**<br>**Gaussian fPET** | **Rest** | 5.77 | 30.23 | 25.07 |
|  | **Win** | 3.68 | 43.77 | 37.41 |
|  | **Loss** | 3.82 | 35.49 | 25.34 |

A comparable pattern emerged for the 2 s edNLM fPET data (see Tables 1 & 2). Good reliability was observed at rest in the NAcc (ICC = 0.66; CV = 3.30). Fair reliability was again found during the win condition in the caudate (ICC = 0.57; CV = 23.04) and the putamen (ICC = 0.46; CV = 23.75), as well as at rest in the putamen (ICC = 0.54; CV = 3.79). Poor reliability persisted across all regions during the loss condition (ICC = 0.11-0.33; CV = 28.88-40.55) and in the NAcc during the win condition (ICC = 0.29; CV = 35.57). Additionally, poor reliability was observed at rest in the caudate (ICC = 0.39; CV = 5.75).

For 2 s fPET data filtered with an 8mm Gaussian kernel, reliability estimates mirrored those of the 2 s edNLM fPET data (see Tables 1 & 2). Fair reliability was seen at rest in the NAcc (ICC = 0.57; CV = 25.07) and the putamen (ICC = 0.51; CV = 30.23). Further, fair reliability was found during the win condition in the caudate (ICC = 0.54; CV 3.68) and the putamen (ICC = 0.47; CV = 43.77). Poor reliability was consistently observed across all regions during the loss condition (ICC = 0.08-0.34; CV = 3.82-35.49), in the caudate at rest (ICC = 0.38; CV = 5.78) and in the NAcc during the win condition (ICC = 0.27; CV = 37.41).

### BOLD-derived neuronal activation

For fMRI beta values (see Tables 1 & 2), good reliability was noted during the feedback win condition in the NAcc (ICC = 0.65; CV = 88.17) and during the feedback loss condition in the NAcc (ICC = 0.67; CV = 244.48) and caudate (ICC = 0.61; CV = 134.15). While fair reliability was found in the putamen across all conditions (ICC = 0.40-0.45; CV = 46.48-379.33), in the caudate during the feedback win condition (ICC = 0.56; CV = -143.33) and cue win condition (ICC = 0.41; CV = 76.70), as well as in the NACC during the cue win condition (ICC = 0.46; CV = 75.76) and the cue loss condition (ICC = 0.46; CV = 82.74). Poor reliability was observed in the caudate during the cue loss condition (ICC = 0.26; CV = 77.22).

## Discussion

This study examined the test-retest reliability of task-induced changes in simultaneously acquired dopamine synthesis measured with 6-[^18^F]FDOPA fPET and BOLD fMRI during the MID task in healthy participants. Our findings reveal region-, condition-, and processing-dependent variability in test-retest measures. Fair to good reliability was observed for task-induced dopamine synthesis in the NAcc and putamen during rest, as well as in the caudate and putamen during the win condition, whereas the reliability of task-induced dopamine synthesis during the loss condition was consistently poor. On the other hand, task-induced betas derived from BOLD fMRI showed good reliability in the NAcc during the feedback conditions and in the caudate during the feedback loss condition. Fair reliability was observed in all other regions across all conditions, apart from poor reliability, which was found in the caudate during the cue loss condition. These findings highlight the challenges of assessing reward-related circuitry and dopaminergic function during MID task performance but also emphasize the complementary nature of fMRI and fPET (34).

This complementarity arises from the fundamental differences between the two modalities. 6-[^18^F]FDOPA fPET measures dopamine synthesis at the molecular level (16,35,36), distinguishing it from the composite and indirect fMRI BOLD signal, which reflects hemodynamic changes, blood oxygenation, and vascular factors (11,12,37). Yet, this specificity is reliant on ionising radiation as well as a more complex and expensive acquisition and analysis pipeline. Contrary to fPET, BOLD fMRI offers excellent spatial and temporal resolution (11,12,38). However, it is more susceptible to motion-artifacts, physiological noise, and scanner-related instabilities (13,14). Furthermore, traditional 6-[^18^F]FDOPA fPET captures data in 30 s frames in a block design, potentially limiting the sensitivity to rapid changes in dopamine synthesis (15). In our analysis we increased the temporal resolution of 6-[^18^F]FDOPA fPET to 2 s based on previous work on high temporal resolution fPET with [^18^F]FDG (23). A key finding of this study is the slight difference in reliability between the 30 s and 2 s edNLM-filtered fPET data. This indicates that 6-[^18^F]FDOPA fPET retains robust reproducibility even at high temporal resolution, without a significant loss of signal quality or an increase in noise. This capability represents a significant methodological advantage of fPET over classical PET imaging approaches, which typically require longer acquisition windows (35,36). The ability to resolve dopamine dynamics on the scale of seconds opens new avenues for studying fast task-related changes in dopaminergic function while maintaining quantitative reliability. This observation will enable the examination of dopamine synthesis in an event related manner in the future enabling to investigate dopamine dynamics during different phases of reward processing. However, our high temporal resolution 2 s fPET frames were also analysed in a block design to allow a direct comparison between 30 s and 2 s fPET data after edNLM filtering. fMRI on the other hand was analysed in an event related manner allowing to distinguish between the anticipation and outcome phases of reward processing (31). Our findings suggest that 6-[^18^F]FDOPA fPET can achieve reliability comparable to fMRI in reward-related regions (e.g., caudate) while offering more direct insight into the underlying neurotransmitter dynamics.

Test–retest reliability of 30 s and 2 s 6-[^18^F]FDOPA fPET data processed with edNLM filtering was comparable to that of [^18^F]FDG fPET during a cognitive task (25). However, [^18^F]FDG fPET acquired at rest showed higher reliability than both 30 s and 2 s 6-[^18^F]FDOPA fPET data (25). For fMRI, our reliability estimates were consistent with prior studies using the MID task with larger samples (39), as well as with test-retest findings from other task-based fMRI paradigms such as finger tapping (40) and decision making under risk (41).

### Comparison of edNLM vs. Gaussian Kernel Filtering

Furthermore, we compared the reliability of different filtering strategies. We previously reported that edNLM filtering enhances reliability for task-specific fPET (26). In our current analysis, data filtered with edNLM yielded similar reliability when compared to filtering with a traditional 8mm Gaussian kernel. The gaussian smoothing is a computationally efficient, though less sophisticated denoising method while edNLM filtering offers a more advanced and adaptive approach but requires longer processing time. The comparable reliability of both filtering strategies suggests that for high temporal resolution 6-[^18^F]FDOPA fPET the choice of filtering method may be guided by computational resources and research goals rather than by absolute performance. Here the 8mm Gaussian kernel might offer a simple yet effective solution for large-scale studies or where processing speed is critical.

### Limitations and Future Directions

Our findings should be interpreted considering certain limitations. Our sample size of 25 healthy subject was limited, which may have decreased the observed reliability, particularly for the more variable task conditions. Furthermore, inter subject variability must be considered. Subjects were measured approximately 8 weeks apart. Previous literature suggests that multiple variables which may change over the course of days or weeks like sleep quality, affective state and stress influence reward-related behaviour (42).

Furthermore, a potential practice effect must be considered, since subjects completed the MID task twice. The first time the reaction to the novel task might have elicited a stronger or different neural response, potentially reducing the reproducibility of our analysis. However, a previous fMRI study using the MID task found no improvement in MID task performance across sessions in healthy subjects, contrary to this prediction (43).

Additionally, it is important to note that 6-[^18^F]FDOPA fPET Ki values were analysed using a block design rather than an event-related approach. Event-related analysis of the 30-second frames is not possible because the different phases of reward processing during the MID task, such as the anticipation and outcome phases, are much shorter. Nevertheless, since the 2 s frames show reliability comparable to that of the 30 s frames, future advancements may allow to assess these distinct phases also with fPET in an event-related manner.

## Conclusion

Our study demonstrates moderate region- and condition dependent test-retest reliability of 6-[^18^F]FDOPA fPET and BOLD fMRI during MID task performance. Neither BOLD fMRI nor 6-[^18^F]FDOPA fPET showed a clear superior test-retest reliability across all regions of interest (caudate, putamen and NAcc). However, 6-[^18^F]FDOPA fPET offers molecular specificity for dopamine synthesis (15,35,36), whereas BOLD fMRI measures a composite signal (11,12). Notably, 6-[^18^F]FDOPA fPET retained comparable reliability also for high temporal resolution 2s frames suggesting the possibility of further event-related analysis and direct comparison to BOLD fMRI at a similar temporal resolution (11). Thus, our findings highlight the complementary nature of BOLD fMRI and 6-[^18^F]FDOPA fPET for investigating reward processing and underscore the chance of using high temporal resolution 6-[^18^F]FDOPA fPET to study dopamine synthesis dynamics even closer.

## Acknowledgments

We thank the graduated team members and the diploma students of the Neuroimaging Lab (NIL, headed by R. Lanzenberger) as well as the clinical colleagues from the Department of Psychiatry and Psychotherapy for clinical and/or administrative support. G. Schlosser, E. Briem, A. Mayerweg, L. Artmeier and S. Klug were supported by the MDPhD Excellence Program of the Medical University of Vienna. M.Reed and C. Milz are recipients of a DOC Fellowship of the Austrian Academy of Sciences at the Department of Psychiatry and Psychotherapy, Medical University of Vienna.

## Author contribution statement

R.L., M.B.R, A.H., P.A.H, M.H. designed the study. G.S., P.A.H., S.G., S.K., B.E., E.B., A.M., L.A., G.M.G., C.S., M.M., C.M., P.F., L.N., S.R. acquired the data. M.B.R. performed data preprocessing and analysis. M.B.R and G.S. interpreted the results. G.S. drafted the manuscript. All authors reviewed, edited, and approved the final manuscript.

## Ethical considerations

The study was approved by the Ethics Committee of the Medical University of Vienna (ethics number: 2321/2019) and all procedures were carried out in accordance with the Declaration of Helsinki.

## Consent to participate

After detailed explanation of the study protocol, all subjects gave written informed consent. All subjects were insured and reimbursed for their participation.

## Consent for publication

Not applicable.

## Data availability

Raw data will not be publicly available due to reasons of data protection. Processed data and custom code can be obtained from the corresponding author with a data sharing agreement, approved by the departments of legal affairs and data clearing of the Medical University of Vienna.

